# Cleavage of the vascular matrix attracts glioblastoma cells to infiltrate the brain parenchyma

**DOI:** 10.1101/2024.08.02.606279

**Authors:** Lara Perryman, Anette M Høye, Thomas R Cox, Ann-Marie Baker, Jan Strøbech, Lidia Leonte, Lukram B Singh, Sergey Popov, Lasse G Lorentzen, Raphael Reuten, Hans Skovgaard Poulsen, Michael J Davies, Chris Jones, Janine T Erler

## Abstract

**Background:** Glioblastoma is a highly aggressive brain cancer and, unlike many other cancers types, the median survival for patients after treatment (14.6 months) has barely improved in the last 20 years. Infiltrative growth into the surrounding brain parenchyma facilitates tumor recurrence and ultimately the death of the patient - novel therapies targeting this process are desperately needed. Lysyl oxidase inhibition has been shown to decrease invasive growth in a variety of solid tumours and is a potential therapy for glioblastoma patients.

**Methods:** Genes highly expressed in the mesenchymal subtype of glioblastoma were analyzed in a data set from the Cancer Genome Atlas and tissue microarrays. Two patient-derived human glioblastoma stem cell lines were used to assess the involvement of lysyl oxidase (LOX). The effect of LOX on infiltration was examined in an organotypic brain slice assay and in an orthotopic mouse model. Chemotactic assays, protease and cleavage arrays were used to assess the underlying mechanism behind LOX-mediated infiltration. The orthotopic model was used to evaluate potential clinical utility of targeting LOX in glioblastoma.

**Results:** LOX is overexpressed in the mesenchymal glioblastoma subtype and strongly associated with poor patient survival. LOX expression upregulates MMP7 expression, which subsequently cleaves the vascular matrix resulting in increased chemotaxis of glioblastoma cells.

**Conclusions:** We have uncovered a novel mechanism of glioblastoma infiltration and suggest that targeting LOX represent an effective therapeutic approach blocking glioblastoma infiltration.

**Importance of the study:** The ability of glioblastoma cells to infiltrate the surrounding normal brain tissue facilitates their evasion of current therapies, leading to tumor recurrence and ultimately the death of the patient. To improve targeted therapies for glioblastoma patients we need to understand the molecular mechanisms of glioblastoma cell infiltration and how cells interact with the unique microenvironment of the brain. We have identified a novel mechanism whereby tumor-derived LOX mediates chemotaxis of glioblastoma cells to the laminin rich perivascular niche, enabling infiltrative growth. Inhibiting this infiltrative pathway is a potential anti-invasive therapy that is desperately needed for glioblastoma patients.

## Introduction

Infiltrative growth in glioblastoma facilitates tumor recurrence and is ultimately responsible for the death of patients. Current methodologies, such as MRI and surgical techniques, are unable to visualize infiltrative cells that escape into the brain parenchyma from resection margins^1^. It has been suggested that infiltrative cells have a lower proliferative index, which enables them to be more resistant to radiation and chemotherapy^2^. It is vital to understand the mechanisms and dynamics of cell infiltration to identify novel anti-invasion targeted therapies in order to treat recurrent growth in glioblastoma.

A major part of the normal brain parenchyma consists of extracellular matrix (ECM). The ECM can broadly be divided into two distinct regions, which is the aforementioned parenchyma and the basement membrane (BM). The ECM in the brain parenchyma is unique since it lacks structural proteins like laminins and collagens and is predominantly composed of hyaluronan and proteoglycans with attached chondroitin sulphate and heparin sulphate chains^3^. By contrast, the BM surrounding blood vessels is primarily composed of laminin, collagen and fibronectin^4^. Glioblastoma cells are known to preferentially invade the area surrounding the blood vessels, which has been coined the “perivascular niche” (PVN)^5,6^. It has been shown that the cells in the PVN have ‘stem-like’ properties, resulting in increased resistance to radiation and chemotherapy^7–9^.

Laminins are a major ECM component of the PVN. They form a sheath-like network surrounding blood vessels. It has been shown that astrocytes express laminin-111 and -211 while endothelial cells express laminin-411 and -511^10^. Laminins have been shown to increase the migration of neural stem cells^11^. Furthermore, laminin and laminin fragments have been shown to act as chemoattractants for murine Lewis lung carcinomas, breast cancer and human myeloma cells^12–14^.

Lysyl oxidase (LOX) is a secreted copper-dependent amine oxidase, which catalyzes the crosslinking of collagens and elastin in the ECM contributing to an increase in tensile strength and integrity of tissues^15^. Over expression of LOX is associated with fibrotic diseases including NASH, system sclerosis and is thought to drive invasion and metastasis in solid tumor types, such as breast, prostate and head and neck^15^. Pan-LOX inhibitor beta-aminopropionitrile (BAPN) has been utilized has been shown to inhibitor invasion^16^ and tumour growth in PTEN null models of glioblastoma proposing a mechanism of LOX as a chemoattractant to macrophages^17^.

LOX and matrix metalloproteinase-7 (MMP-7) were included in the gene signature for the mesenchymal subtype in the latest classification of glioblastoma subtypes^18^. Here, we confirm LOX being highly expressed in mesenchymal glioblastoma and uncover a novel mechanism whereby LOX up-regulates MMP-7, which cleaves laminin, promoting chemotaxis and thereby infiltration of glioblastoma cells into the brain parenchyma.

## Materials and Methods

### Patient sample analysis

We analyzed gene expression in the Cancer Genome Atlas (TCGA) glioblastoma patient data set previously published (http://tcga-data.nci.nih.gov/docs/publications/gbm_exp/), TCGA_unified_CORE_ClaNC840.txt)^17^. The median gene expression was calculated for each gene from all samples designated to the Mesenchymal subtype. The genes were then ordered by highest to lowest median gene expression. We additionally performed immunohistochemical (IHC) staining of LOX on a patient tissue microarray (TMA) of adult glioblastoma (n=165) as described^18,19^. LOX staining of tumors was assessed by a pathologist and graded based on the intensity of staining, where 0 is negative, 1 is the lowest, 2 is intermediate and 3 is the highest positive staining.

The Ivy Glioblastoma Atlas Project (Ivy GAP) database was accessed on May 2^nd^ 2018 specifically searching for LOX and MMP-7 to evaluate mRNA expression levels.

### Cell lines and culture

The GBM_CPH048 (048) human glioma stem cell line was a kind gift from Hans Skovgaard Poulsen, Copenhagen University Hospital^20^, and the NH101123 (NH10) human glioma stem cell line was a kind gift from Sebastian Brandner, UCL Institute of Neurology^21^. All glioblastoma stem cell lines were cultured in Neural basal media (NB) (Invitrogen) with the following supplements: B27, N2, L-glutamine, Penicillin 0.5%, Streptomycin 0.5%, 20 ng/ml basic fibroblast growth factor (bFGF) and 20 ng/ml epidermal growth factor (EGF; Invitrogen), referred to as NB+. All cell lines were regularly confirmed negative for mycoplasma using a 4-myco polymerase chain reaction (PCR) detection kit (Chembio Ltd, Hertfordshire, UK). We utilized short tandem repeat (STR)-based DNA profiling and multiplex PCR to confirm cell identification (CellCheck, IDEXX laboratories). In addition, cell lines tested negative for murine pathogens by IMPACT testing (IDEXX laboratories).

### Cell manipulations

Sixty percent confluent HEK293T cells were transfected with pCMV6-AC-GFP-LOX lentiviral vectors (Origene) utilizing Lipofectamine 2000 (Invitrogen), as per manufacturer’s instructions. 24 h after transfection, lentiviral supernatants were collected and media replaced. After a further 24 h supernatants were collected again, pooled, filtered, aliquoted and stored at -80°C. The supernatants were used to infect the glioblastoma stem cell lines with the addition of 8 μg/ml of polybrene (Millipore). Stably transfected cells were selected either by puromycin or G418 (Sigma) as appropriate. For *in vivo* monitoring, cells were labelled with a pFU_luc2_eGFP or a Tomato lentiviral vector, a kind gift from Sanjiv Sam Gambhir (Stanford University).

### Western Blot

Cells were plated in a T75 cm^2^ flask (BD Falcon) with or without laminin coating (as described below) and allowed to settle for 48 h. The conditioned medium (CM) and total cell lysate (TCL) were collected and the Western blots were performed as described^18^. Antibodies detecting human LOX (Novus), laminin (Sigma) and MMP-7 (R&D Systems) were used at 1:500.

### Organotypic brain slice invasion assay

As described^22^, brains from female NCR/Nude mice (Taconic) 6-8 weeks, were harvested and embedded in agarose. A vibratome (Leica, VT1200S) was used to cut 300 μm brain sections (Velocity 0.80mm/s, Amplitude 2 mm, Frequency 60 mm/s 1.50 mm), which were then added to membrane inserts (Millicell, Millipore) and incubated at 37°C, 5 % CO_2_ and 95 % humidity, with NB+ medium. The medium was changed every other day. Spheroids of 6000 cells/ 20 μl in 0.6% methylcellulose (Sigma) in NB+ media for 24 h were created, placed on the brain slice and incubated for 7 days before fixing with 4% paraformaldehyde. For immunofluorescence, sections were incubated with a blocking buffer (10% normal goat serum, 2% bovine serum albumin, 0.25% Triton-X (all Sigma) in PBS) for 2 h, washed extensively in PBS with 0.25% Triton-X (PBS-Triton). Sections were incubated with anti-vimentin antibody 1:200 using the same blocking buffer overnight at 4°C, washed with PBS-Triton, and finally incubated with secondary goat-anti mouse Alexa-488 1:1000 and Hoechst 33342 (Sigma) 10 μg/ml in the blocking buffer for 2 h at room temperature. Confocal micrographs were acquired on an Axiovert 200M LSM 520 (Carl Zeiss) using a Zeiss C-Apochromat x10 objective. Images were processed using ZEN 2009 (Carl Zeiss) and ImageJ.

### Orthotopic glioblastoma model

Orthotopic injections were performed in female NCR/Nude mice (Taconic) 6-8 weeks old anaesthetized with isoflurane. An incision through the skin was made along the midline of the skull, and a burr hole drilled 1 mm lateral and 2 mm anterior to the bregma and 3 mm deep. The mice received stereotactic intracranial injections of 1-2x10^5^ cells. *In vivo* monitoring of growth was performed by anaesthetizing the mice and intraperitoneal injection of luciferin (120 mg/kg) (Caliper Lifesciences) and imaged using an IVIS (PerkinElmer). In survival experiments, mice were euthanized based on short-term weight loss and condition. Dosing of tumors began at 73 days, as tumors were observable via bioluminescent signal. Mice were randomly assigned into treatment and control groups. Treatments included: daily intraperitoneal injection of 100 mg/kg β-aminopropionitrile (BAPN) or saline, and LOX antibody (Open Biosystems)^23^ or rabbit IgG control (Sigma) given intraperitoneal twice weekly at 1mg/kg. All animal procedures were completed in accordance with the Danish Health Ministry guidelines for animal experimentation.

### Conditioned Medium (CM)

CM was prepared by seeding 1-2x10^6^ cells in neural basal media without supplements (NB0) or NB+ media for 24 h. The media was collected, filtered in 0.45 μm filter (Sartorius) and, for Western blot analysis, concentrated by centrifugation (3000g for 20 min) using a 10 kDa molecular mass cutoff (MWCO) column (Sartorius).

### Boyden chamber chemotactic assay

A modified Boyden chamber was utilized^11^ to evaluate chemotaxis. Twenty-four well plates were coated with laminin-111 (10 μg/ml, Sigma), collagen I (10 μg/ml, Invitrogen), fibronectin, or poly-lysine (100 μg/ml, Sigma) as negative control, and incubated for 2 h at 37°C, washed with PBS before media with treatments were added. Uncoated inserts (8um pores, BD Biosciences) with 1x10^4^ cells were added to wells. Cells adhering to coated wells were counted after three days to determine the number of cells undergoing chemotaxis. In some experiments CM was pre-incubated with different lower well coatings for 24 h and then transferred to a new well before cells were added. The well containing the coating modified by CM pre-incubation was washed and new NB+ media added. Recombinant proteins, conditioned media and inhibitors were added to wells at concentrations described below. Recombinant (r) proteins rLOX (Origene), active rMMP-7 (Millipore) and rPro-MMP-7 (R&D systems) were used at 0.5 μg/ml. Rabbit anti-human LOX polyclonal antibody (Open Biosystems)^23^ and rabbit IgG (Sigma) were used at 5 μg/ml, and BAPN (Sigma) was added at either 20 μM or 200 μM as indicated.

### Protease array

The presence of 34 human proteases in CM was investigated using a Human Protease Array Kit (R&D Systems) according to manufacturer’s instructions. Briefly, the membranes were hydrated, and CM was mixed with a cocktail of biotinylated detection antibodies and incubated with the array overnight. For detection, membranes were incubated with streptavidin-HRP followed by an HRP substrate and developed using autoradiography film (GE Healthcare). Images were analyzed with ImageJ (National Institutes of Health).

### Cleavage assay

Six-well plates were coated with 1 ml 100 μg/ml Laminin (Sigma) as described above, then treated with 1 ml of 4 or 8 μg/ml active rMMP-7 (Millipore) in PBS or PBS only and incubated at 37°C for 48 h. Fragments were detected in filtered and concentrated (using 30 kDa MWCO spin filters, Sartorius) supernatants. Laemlli sample buffer was added to samples which were boiled and run on 4-12% Bis-Tris Nupage SDS-PAGE gels (Invitrogen) and visualized by silver staining (Thermo Scientific) following the manufacturer’s protocol.

### Statistics

Results are presented as mean ± s.e.m. Statistical analysis was performed using two-tailed unpaired Student’s t-test or one-way analysis of variance (ANOVA). Kaplan-Meyer survival curves were analyzed by log-rank Mantel-Cox test. Statistical analysis was performed using GraphPad Prism.

## Results

### LOX expression is up-regulated in the mesenchymal subtype of glioblastoma

It has been previously shown that the mesenchymal subtype of glioblastoma is associated with poor survival^26^ and that LOX is a part of the mesenchymal gene signature^18^. The mesenchymal subtype is also the most necrotic, which induces a hypoxic environment, which is known to up regulated LOX expression.^25^} Analysis of a published gene expression microarray dataset^19^ from glioblastoma patients revealed *LOX* as the gene most highly up-regulated in the mesenchymal subtype (Figure 1A). In the original classification of glioblastoma subtypes^19^ LOX mRNA expression was significantly higher in the mesenchymal subgroup compared to the other glioblastoma subtypes proneural, neural, classical and C-GIMP (Figure 1B). Importantly, patients with high LOX mRNA expression has a significantly worse overall survival compared to patients with low LOX (n=150) (Figure 1C). Moreover, immunohistochemical staining for LOX expression on a lioblastoma patient TMA (n=165) found that 76.4% of tumor cores were positive for LOX (scoring grade 1-3) (Figure 1D-E). However, infiltrative growth was not assessed in the tissue microarray.

**Figure 1.**
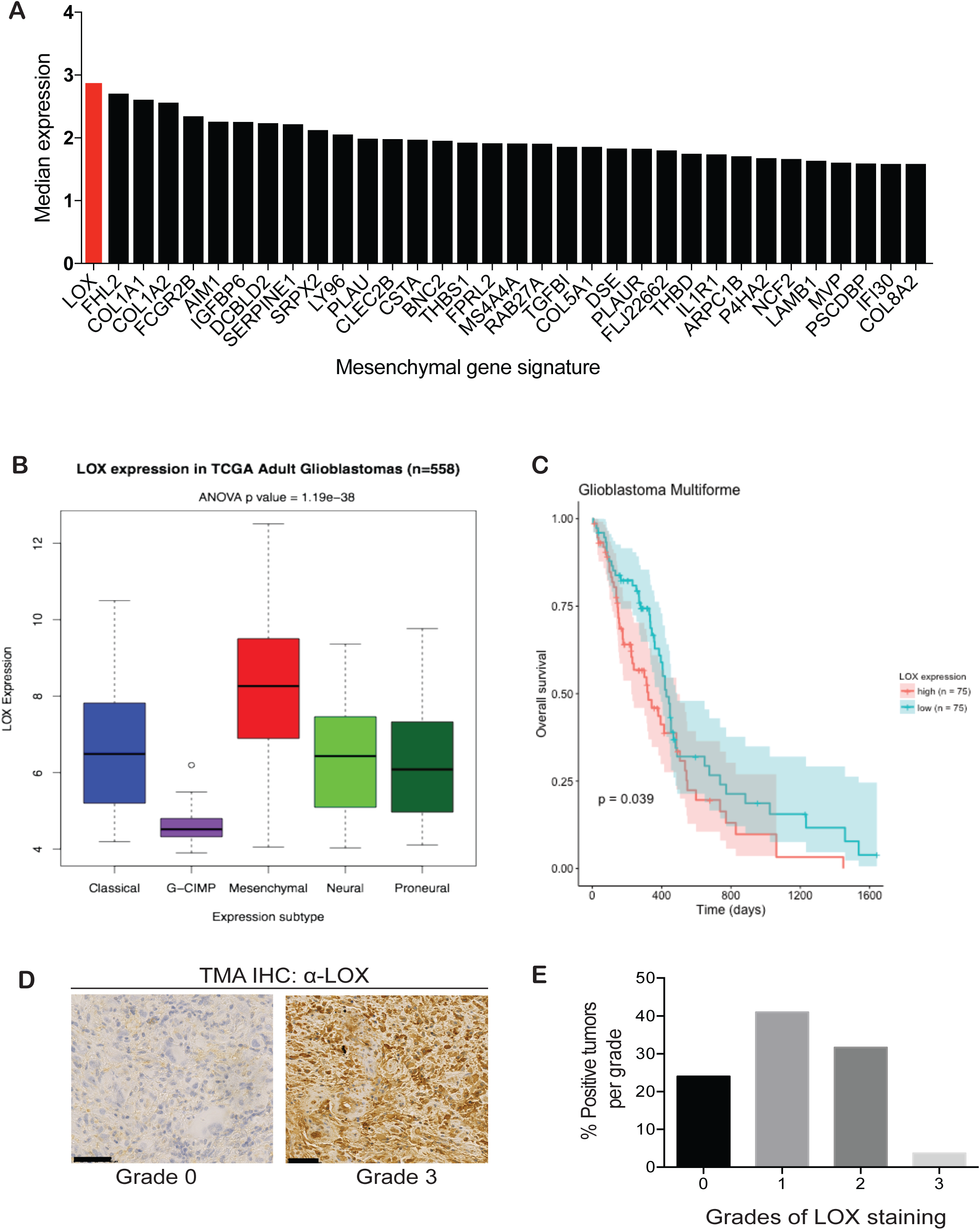
LOX expression is associated with the Mesenchymal subtype of glioblastoma. A) The median gene expression levels of the most highly upregulated genes in the Mesenchymal subtype of glioblastoma in a TCGA gene expression data set (N=558)^17^. B) LOX gene expression is highest in the Mesenchymal subtype (data are analyzed by ANOVA, P=1.19e-38, N=558) compared to classical, G-CMP, neural and the proneural subtype in adult glioblastoma. C) Overall survival of patients with high and low LOX expression in a TCGA glioblastoma dataset (P=0.039, N=150) D) Immunohistochemistry (IHC) staining of LOX in a glioblastoma patient tissue microarray (TMA). Left, example of a negative LOX staining (Grade 0). Right, example of a LOX positive staining (Grade 3). E) Scoring of TMA in D). 76.4% of patient samples are positive for LOX expression (N=165).

### LOX affects glioblastoma invasion *ex vivo* and *in vivo*

We consider that glioblastoma patient survival is intimately related to the infiltrative phenotype of the tumor. As LOX expression has been shown to drive invasion in a variety of other cancer types^15^ we hypothesized that LOX can affect infiltration in glioblastoma. To evaluate this, we took advantage of two human patient derived glioblastoma stem cell lines that express different levels of LOX. NH10 expresses LOX while 048 expresses negligible amounts of LOX (Figure 2A). The 048 cells were either transduced with an empty vector (048EV) or a LOX containing vector (048LOX) using lentiviral infection (Figure 2B, Supplementary Figure 1A-B) to evaluate whether expression of LOX affects glioblastoma infiltration.

**Figure 2.**
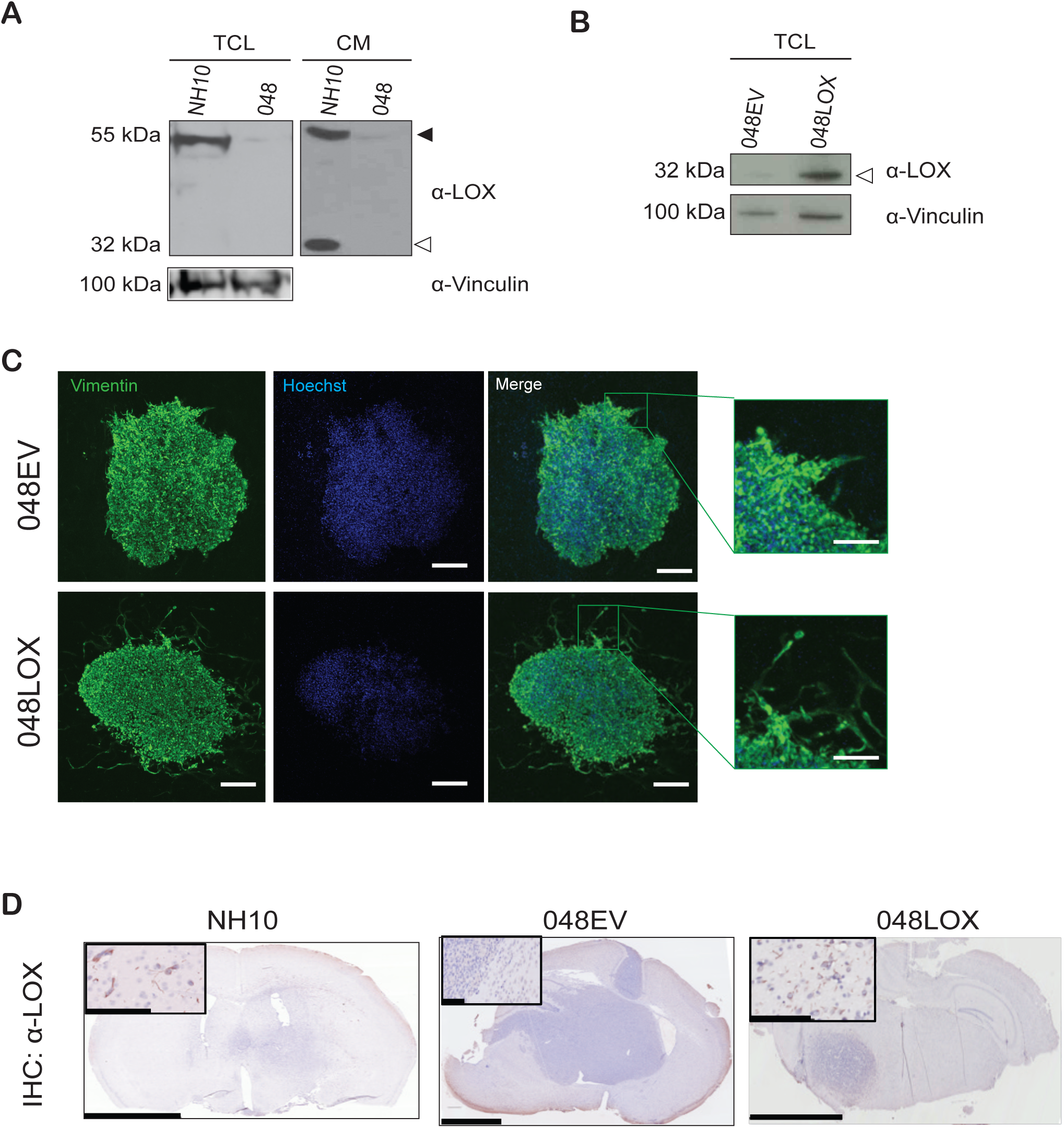
LOX expression is associated with infiltrative glioblastoma *in vivo*. A) Western blot of total cell lysate (TCL) and conditioned medium (CM) from NH10 and 048 cell lines. Black arrowhead denotes the proform of LOX. Open arrowhead denotes the active form of LOX. B) Western blot of TCL from 048 empty vector (048EV) and 048 overexpressing LOX (048LOX) cell lines. C) An organotypic brain slice assay with 048EV or 048LOX spheroids incubated for 7 days. Shown are representative maximum-projections of Z-stacks of confocal micrographs. The slices are stained for vimentin (a human specific antibody) and Hoechst (nuclear stain). Scale bars are 200μm. Inserts show cells at the sphere boundary, scale bars 100μm. D) Representative LOX IHC staining of orthotopic patient derived tumors. Left, an NH10 orthotopic xenograft which is highly invasive and the infiltrative cells are positive for LOX expression. Middle, an 048EV non-infiltrative glioblastoma intracranial tumor which does not express LOX. Right, 048LOX orthotopic tumor presents with infiltrative cells. Scale bars are 2.5mm. Insert highlights LOX staining at the invasive edge of the tumor, scale bars are 100μm.

We employed an *ex vivo* organotypic brain slice assay to study the invasion of the modulated cell lines. Using this assay, the 048EV spheroids did not show infiltrative behavior. In contrast, 048LOX spheroids reveal extensive infiltration both as tube-like structures and as diffuse single cells (Figure 2C).

Moreover, 048LOX cells proliferated significantly less compared to 048EV cells (Supplementary Figure 1C). These findings strongly suggest that LOX expression causes a switch from a proliferative to invasive phenotype driving glioblastoma infiltration.

### The expression of LOX facilitates chemotaxis of glioblastoma cells towards laminin

We have previously shown that LOX modification of premetastatic sites recruits CD11b+ cells which produce MMP-2 and thereby modifies the BM by releasing chemotactic fragments^25^. Therefore, we hypothesized that directional cell movement towards soluble or solid-state gradients, chemotaxis, could initiate the infiltrative behavior observed by LOX expression. In order to investigate this, the wells of a Boyden chamber (not the insert) were first coated with poly-lysine (Poly_L), collagen I (Col I), plasma fibronectin (FN) or laminin-111 (Lm-111), then NB+ media was added to the wells and tumor cells added to the insert (Figure 3A). Tumor cells rarely attached to the top or bottom of the insert, but cells were instead found to attach to the coated wells; these were counted as ‘chemoattracted’ cells. Chemotaxis of the glioblastoma cells was not observed in wells coated with poly-lysine, fibronectin or collagen I (Figure 3B). Only cells expressing LOX (048LOX and NH10) were significantly more attracted to laminin-111 coated wells compared to 048EV (Figure 3B). The chemotaxis model was not specific to laminin-111, as other laminin trimer coatings also induced chemotaxis of NH10 and 048LOX, but not 048EV (Supplementary Figure 2A). Two LOX inhibitors, BAPN, a LOX family small molecule inhibitor, and a function-blocking LOX antibody significantly decreased LOX-specific chemotaxis to laminin (Figure 3C). These results indicate that LOX activity facilitates glioblastoma cell chemotaxis towards the laminin matrix.

**Figure 3.**
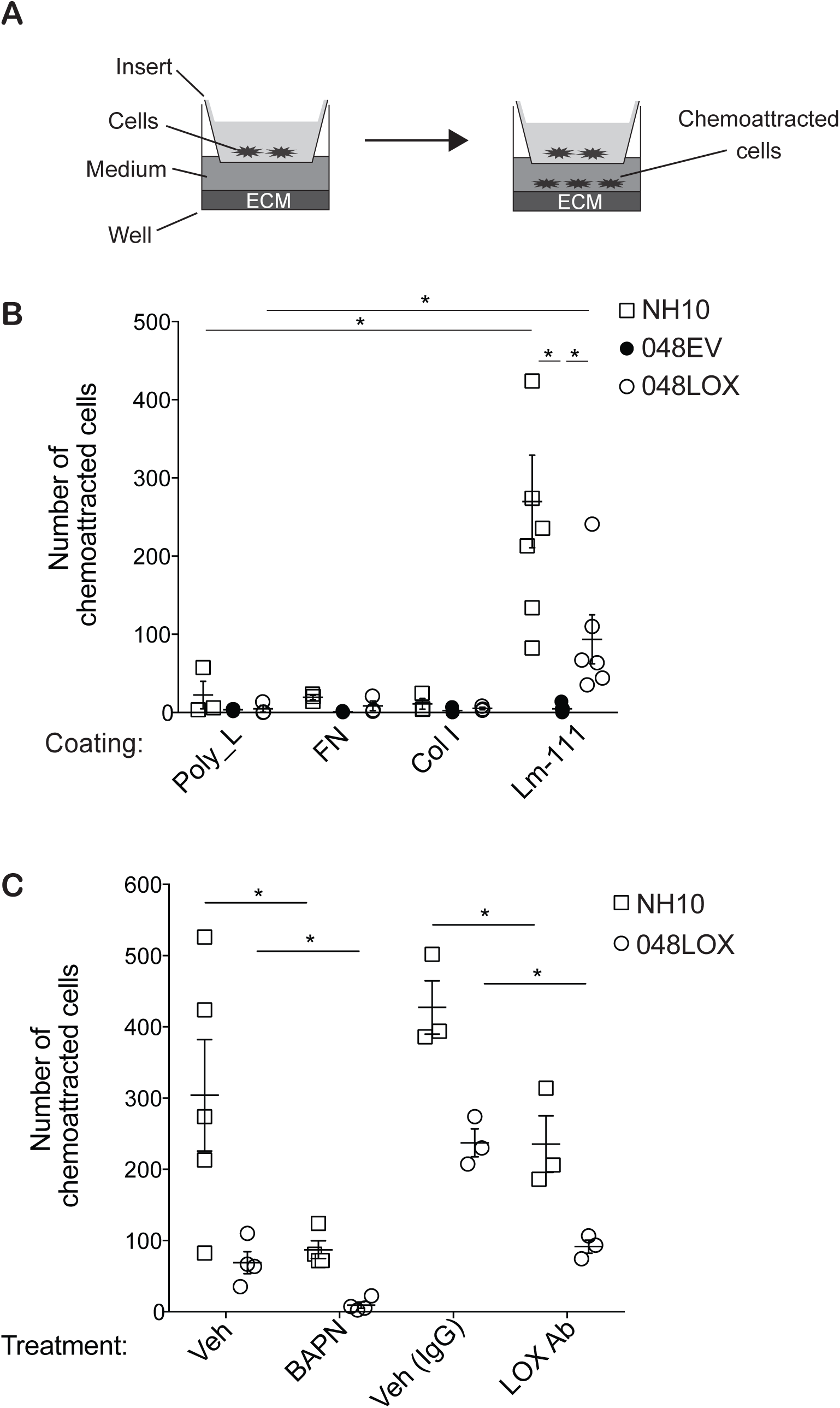
LOX expression facilitates chemotaxis of glioblastoma cells towards laminin. A) Schematic of the Boyden chamber chemotaxis assay as described in Materials and Methods. B) The LOX expressing lines NH10 and 048LOX undergo significantly more chemotaxis when a laminin-111 (Lm-111) coating is present compared to poly-lysine (Poly_L) coated negative control, but not by fibronectin (FN), or collagen I (Col I). NH10 and 048LOX also significantly underwent chemotaxis toward laminin-111 compared to 048EV. Chemotaxis of 048EV cells is not induced by any coatings. (*= p<0.05, n≥3) C) BAPN and the LOX antibody (LOX Ab) decrease the chemotaxis of NH0 and 048LOX to laminin-111 compared to no treatment (NB+ alone). (*= p<0.05, n=5). Data are means ± s.e.m. using Student’s *t-*test.

To test whether LOX had a direct or indirect effect on the chemotaxis of glioblastoma cells towards laminin, CM from control cells (048EV), LOX expressing cells (048LOX or NH10) or recombinant LOX (rLOX) was added to Boyden chamber wells coated with laminin (Figure 4A). Complete neurobasal medium (NB+), not incubated with cells, was included as a negative control. Only CMs from 048LOX or NH10 induced significantly more chemotaxis of 048EV cells (Figure 4B). rLOX on its own did not have an effect on the chemotaxis of 048EV cells, demonstrating that LOX does not have a direct effect on chemotaxis (Figure 4B).

**Figure 4.**
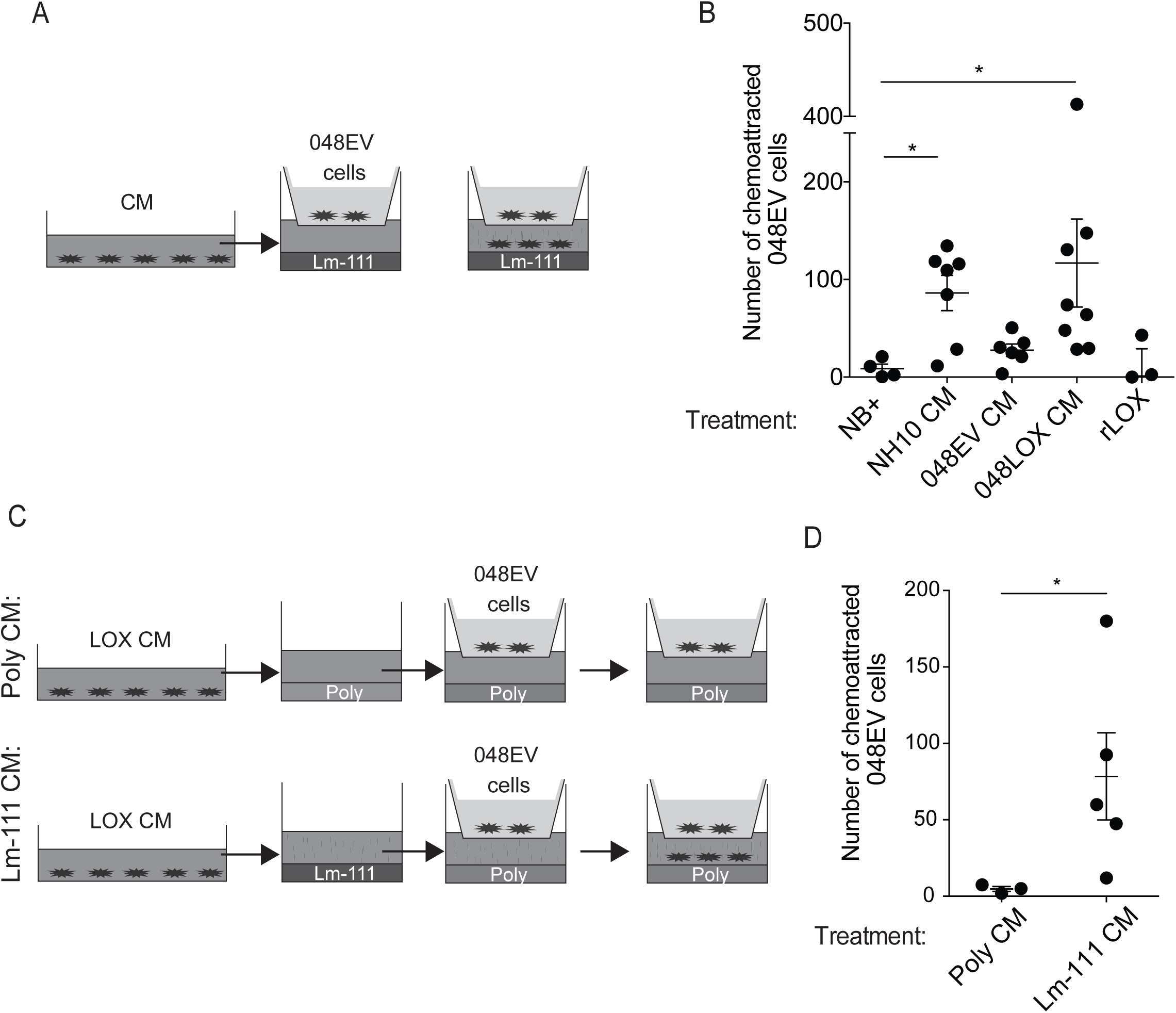
Conditioned medium from LOX expressing cells generates laminin fragments driving chemotaxis. A) Schematic of the Boyden chamber chemotaxis assay with conditioned medium (CM) generation as described in Materials and Methods. B) 048EV cells treated with CM from LOX expressing cells (NH10 and 048LOX) undergo significantly more chemotaxis toward the laminin-111 coating compared to NB+ (negative control), 048EV CM and recombinant LOX (rLOX) only. (*= p<0.05, n≥3). C) Schematic of first generating Poly lysine CM (Poly CM) and laminin-111 CM (Lm-111 CM) and next applying them to a Poly lysine (Poly) coated well for the chemotaxis assay. D) Only Lm-111 CM, and not Poly CM, caused chemotaxis of 048EV cells to poly-lysine coated wells. (*= p<0.05, n≥3). Data are means ± s.e.m. using ANOVA with Kruskal-Wallis test (B) or Student’s *t-*test (D).

As rLOX did not have a direct chemotactic effect, we investigated whether this was caused by fragments from the laminin coating, or if a factor in the CM from LOX expressing cells induced chemotaxis. We generated 048LOX CM and incubated this for 24 h with an ECM coating, either poly-lysine (Poly CM) as negative control or laminin (Lm-111 CM) (Figure 4C). These CMs were transferred to a new well coated with poly-lysine, and 048EV cells were added to the inserts (Figure 4C). Significant chemotaxis only occurred when Lm-111 CM was applied to the wells (Figure 4D). Poly CM or 048LOX CM on a poly-lysine coating did not stimulate chemotaxis (Figure 4D). We also showed that the coating remaining after the generation of 048LOX CM did not cause chemotaxis of 048EV cells (Supplementary Figure 2B). These data suggest that factors in the LOX CM interacts with the laminin coat releasing soluble chemoattractant laminin fragments.

### LOX upregulates MMP-7 expression driving laminin-mediated chemoattraction

MMPs can cleave ECM components to create functional fragments that can induce or modulate cell behavior, including chemotaxis^28^. In order to identify whether a protease was responsible for the observed chemotaxis of glioblastoma cells towards laminin, we examined CMs from 048EV and 048LOX cells using a protease array (Figure 5A). The proteases that were differentially upregulated in the 048LOX CM compared to 048EV CM included cathepsins and a number of MMPs, including the known laminin-332 proteases MMP-7 and MMP-12 (Figure 5A-B). MMP-12 was excluded as the responsible MMP, since recombinant MMP-12 (rMMP-12) and an MMP-12 inhibitor did not affect chemotaxis (Supplementary Figure 2C-D). Interestingly, the mRNA expression level of MMP-7 was upregulated in 048LOX cells compared to 048EV (Supplementary Figure 2E). These results are in line with that both LOX and MMP-7 are proposed to be a part of the mesenchymal gene signature in glioblastoma^18^. To corroborate this, we utilized the Ivy Glioblastoma Atlas Project (Ivy GAP) database, which showed that LOX mRNA expression is upregulated in hypoxic regions within the tumor, including: the perinecrotic zone, pseudopalisading cells around necrosis, hyperplastic blood vessels and microvascular proliferation (Figure 5C). Interestingly, the mRNA expression profile of MMP-7 is similar to LOX (Figure 5C).

**Figure 5.**
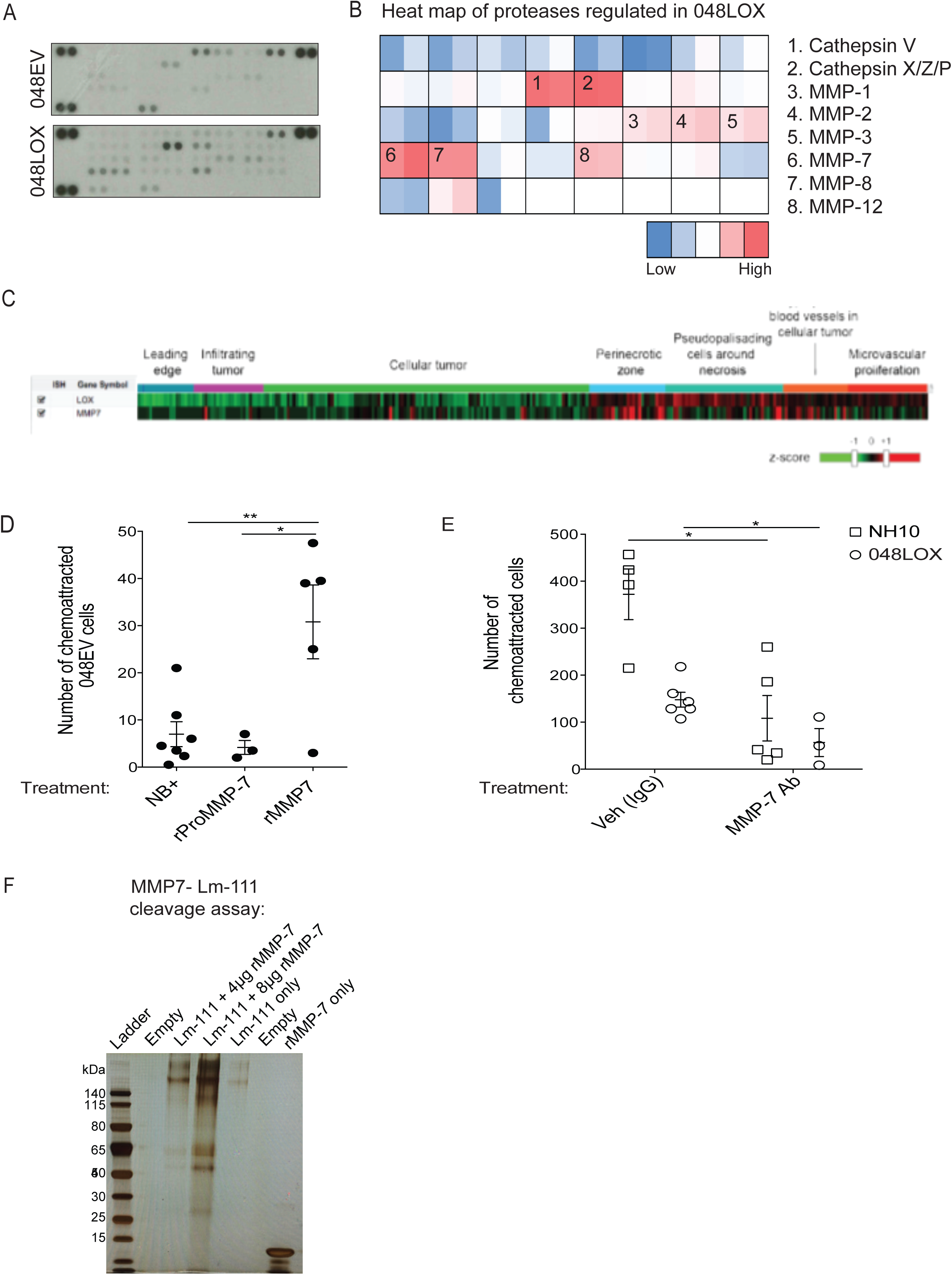
MMP-7 cleaves laminin driving chemotaxis of glioblastoma cells. A) Image of representative protease arrays incubated with CM from 048EV (top) and O48LOX (bottom)(n=2). B) Heat map of the upregulated proteases in 048LOX CM compared to 048EV CM from A). C) RNAseq expression z-score values of LOX and MMP-7 according to the Ivy GAP database. The top horizontal bars represents areas sampled by reference histology (blue: leading edge (n=19), purple: infiltrating tumor (n=24), green: cellular tumor (n=133), light blue: perinecrotic zone (n=26), mint green: pseudopalisading cells around necrosis (n=40), orange: hyperplastic blood vessels in the cellular tumor (n=22), red: microvascular proliferation (n=28)). D) Wells were coated with laminin-111 and 048EV cells were treated with NB+ (negative control), recombinant inactive MMP-7 (rProMMP-7) or recombinant active MMP-7 (rMMP-7). Only active rMMP-7 significantly increases chemotaxis compared to both NB+ and rProMMP-7. (*p<0.05, **p<0.01, N≥3). E) Chemotactic assay towards laminin-111 coated wells where the LOX expressing cells NH10 and O48LOX were treated with control IgG (Veh (IgG)) or an MMP-7 antibody (MMP-7 Ab). Treatment with MMP-7 Ab significantly inhibit chemotaxis of both cell lines compared to IgG. (*=p<0.05, n≥3). F) Silver stained SDS-PAGE gel of supernatants from the MMP7-laminin-111 cleavage assay. The entire concentrated supernatant from each condition was loaded. Wells were coated with laminin-111 and incubated with 4 μg or 8 μg active rMMP-7. As negative controls PBS was incubated with a laminin-111 coated well (Lm-111 only) and an uncoated well incubated with active rMMP-7 only as indicated. Data are means ± s.e.m. using ANOVA with Tukey’s multiple comparisons test (D) or Student’s *t-*test (E).

To determine if the chemotaxis was MMP-7-dependent, active recombinant MMP-7 (rMMP-7) was added to the chemotactic assay with 048EV (low LOX, low MMP-7) cells. Active rMMP-7, and not the recombinant inactive form (rProMMP-7), could drive chemotaxis (Figure 5D). The effects observed due to active rMMP-7, was comparable to those seen with 048LOX CM. Moreover, we were able to block the response in the high LOX lines (048LOX and NH10) by adding an MMP-7 blocking antibody (Figure 5E).

To verify that MMP-7 could generate laminin fragments we completed a cleavage assay utilizing the laminin-111 coat and active rMMP-7. Cleavage of laminin-111 by active rMMP-7 was confirmed by silver staining of the supernatants (Figure 5F). Furthermore, LC-MS/MS identified several peptides in the supernatant where the laminin coat had been incubated with active rMMP7 (Supplementary Figure 3; Supplementary table 1). These findings strongly suggest that LOX drives glioblastoma infiltration by increasing MMP-7 activity to create a chemotactic gradient of laminin fragments.

### LOX is associated with decreased survival in orthotopic xenograft models

Next, we orthotopically injected 048LOX and 048EV cells into the cerebral hemispheres of immunocompromised mice as described above. The survival of animals bearing 048LOX tumors was significantly reduced compared to mice bearing 048EV tumors (Figure 6A). Orthotopic NH10 tumors treated with the LOX inhibitor BAPN or with the LOX-targeting antibody showed a trend towards increased survival, compared to vehicles (Figure 6B-C). In summary these results suggest that targeting LOX may represent an effective new approach to inhibiting glioblastoma infiltration.

**Figure 6.**
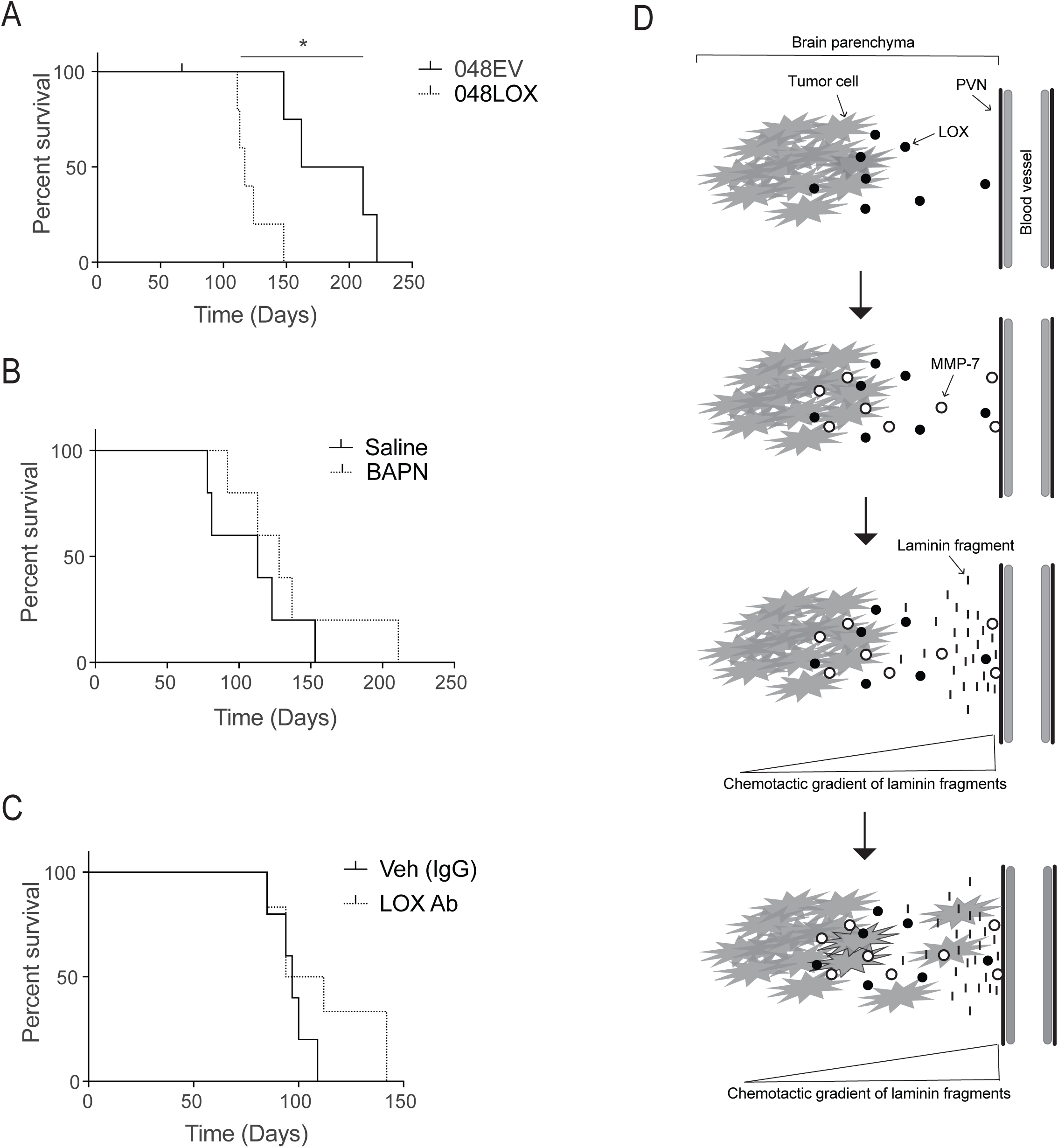
LOX expression affects survival *in vivo*. A) Orthotopic injection of 048EV and 048LOX cells. Survival curves show that 048LOX bearing mice have a lower survival compared to 048EV bearing mice. (*=p<0.05, n=5). B) Orthotopic injection of NH10 cells. Tumors were treated with vehicle (saline) or BAPN (LOX inhibitor). BAPN treated mice have a tendency towards increased survival compared to vehicle treated mice. (p=0.384, n=5). C) Orthotopic NH10 tumors treated with vehicle (IgG) or LOX antibody (LOX Ab). LOX Ab treated mice have a tendency towards increased survival compared to vehicle treated mice (p=0.218, n≥5). Kaplan-Meyer survival curves were analyzed by log-rank Mantel-Cox test. D) Proposed mechanism of action. LOX upregulates MMP-7 expression which then cleaves laminin located in the perivascular niche (PVN). This generates a chemotactic gradient driving the infiltration of glioblastoma cells.

## Discussion

We have investigated potential mediators of glioblastoma infiltration through identifying genes strongly upregulated in the mesenchymal subtype of glioblastoma. We identified LOX as a gene of potential interest, given previous reports showing that LOX is highly up-regulated in several aggressive solid tumor types and since it plays a major role in mediating invasion and metastasis^15^. A relationship between LOX expression and tumor grade in glioblastoma patient samples has been observed previously, but the relationship to infiltrative growth was not investigated^30^.

The role of LOX in glioblastoma infiltration has not been defined previously and the role of LOX expression in infiltrative glioblastoma cells has not been assessed. Orthotopic tumors from the U87MG glioblastoma serum-cultured cell line treated with either oral BAPN or D-penicillamine showed a significant reduction in tumor size^32^. However serum is known to alter the state of glioblastoma cells and result in well-circumscribed, non-invasive tumors *in vivo* that do not recapitulate those observed in patients^33^. *In vitro* migration and invasion studies of serum-cultured cells using Matrigel (where laminin-111 is the main component) treated with BAPN, support our findings that migration and invasion can be reduced by LOX inhibition^33^. BAPN is a small-molecule inhibitor and expected to cross the BBB. However, we speculate that the LOX Ab may have difficulties penetrating an intactBBB and into the brain parenchyma.Therefore, we hypothesize that the LOX Ab reaches areas of the tumor where blood vessels are disorganized, hyperplastic and undergoing microvascular proliferation and is thus able to affect infiltrating cells.

Instead of a direct role, we have shown that LOX upregulates MMP-7 mRNA (Supplementary Figure 2E) and induces the secretion of active MMP-7 protein (Figure 5A-B). We speculate that LOX affects the transcription of MMP-7 as lines of evidence suggest LOX can enter the nucleus and affect chromatin structure and gene transcription by oxidation of nuclear histones^15,34^. Thereafter, laminin-111 is cleaved by active MMP-7 (Figure 5E). In contrast, LOX acts as a direct chemoattractant with mouse embryonic fibroblasts and vascular smooth muscle cells^35^. The enzymatic activity of LOX may oxidize non-active MMP-7 directly, or via the formation of hydrogen peroxide (a product of LOX catalytic activity) which is known to activate MMP-7 via the cysteine switch pathway^36^.

To our knowledge, we have shown for the first time that MMP-7 can cleave laminin-111 (Figure 5E; Supplementary Figure 3; Supplementary Table 1) which then acts as a chemoattractant. Our results also indicate that MMP-7 can cleave a range of laminins (Supplementary Figure 2A), and not only laminin-332. Previously, laminin-111 fragments generated by MMP-2 have been shown to regulate epithelial to mesenchymal transition in embryonic stem cells^29^ complementing our findings.

The PVN, a laminin-rich environment, is central to glioblastoma stem cell maintenance, growth and survival after radiation treatment^37^. Laminins have been shown to be processed by a variety of proteases (cathepsins, elastases, MMPs)^38,39^ to alter BM structure, and drive migration, in a variety of cell types including neutrophils, neural stem cells and cancer cells^11,40–42^. The laminin β1 chain peptide fragments, YIGSR and LGTIPG, have previously been implicated to have an effect on cell attachment and chemotaxis^12,43^. Interestingly, laminin gradients are capable of orientating neurons, astrocytes and neuroblastoma cells^44,46^ providing supporting evidence for our hypothesis that a chemotactic gradient of laminin fragments located in the vascular matrix drives invasion of glioblastoma cells towards the perivascular niche.

In summary, we have identified a novel mechanism of action whereby LOX drives chemoattraction of glioblastoma cells to laminin in the PVN through MMP-7 activity, driving infiltrative growth (Figure 6D). We further show that LOX inhibition can decrease chemotaxis and invasion, and that the expression of LOX has an impact on survival *in vivo*. These findings provide important insight into understanding the mechanisms that drive infiltration in glioblastoma. Moreover, our findings suggest that inhibition of LOX is a potential anti-invasive therapy to increase the survival of glioblastoma patients.

## Acknowledgments

We thank Anna Fossum (BRIC, University of Copenhagen) for her assistance with flow cytometry and Martin Illemann (BRIC, University of Copenhagen) for assistance with immunohistochemistry.

## Authorship

LP and JTE conceived the project. LP, AMH and JTE wrote the manuscript with input from all authors. LP, AMH, TRC, AMB, JS, LL, and LBS designed, performed and analyzed experiments. LGL, RR and MJD designed, performed and analyzed LC-MS/MS samples. SP and CJ analyzed and scored glioblastoma samples. HSP provided reagents.

---

**Supplementary Figure 1.**
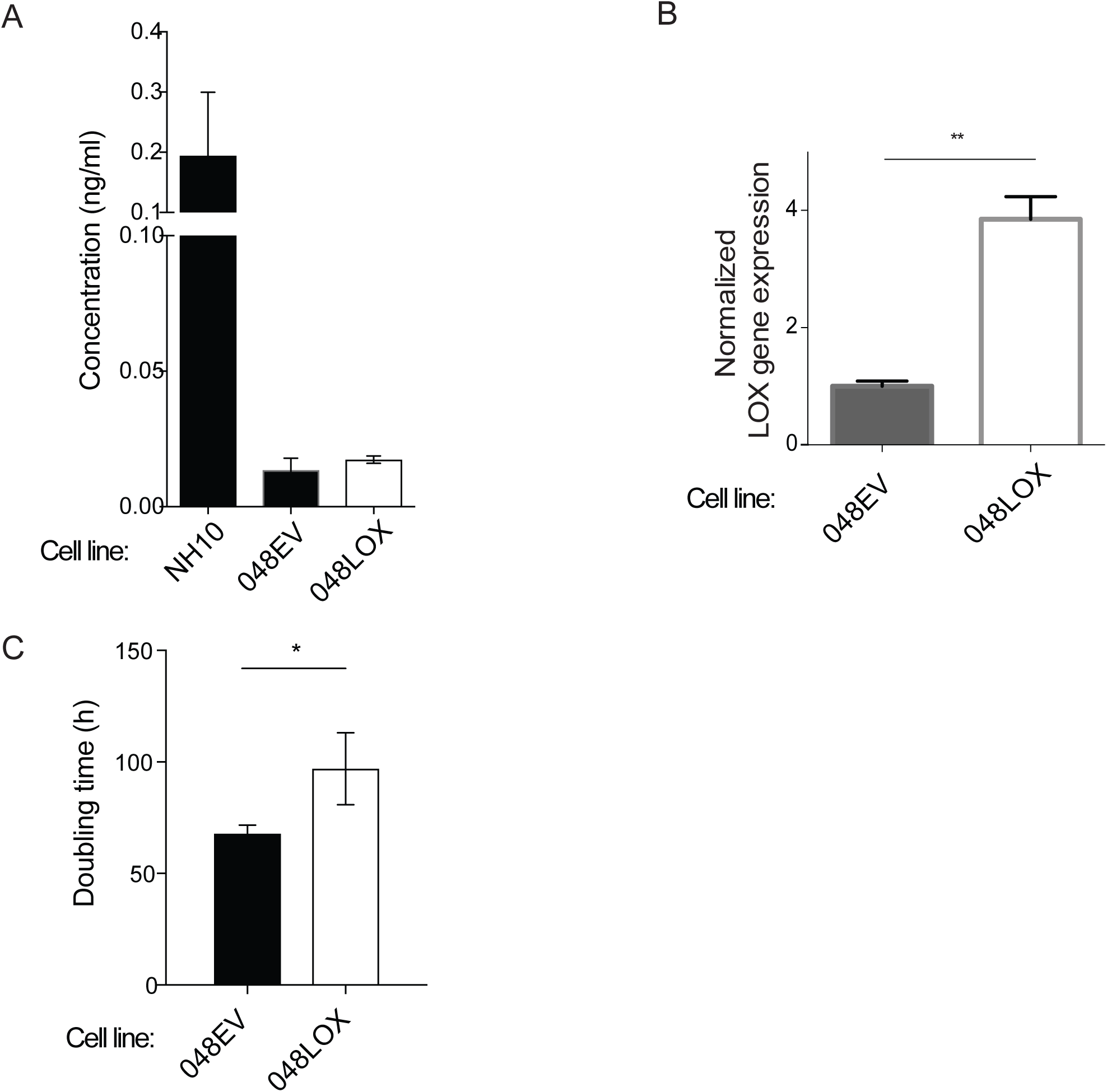
LOX is expressed in invasive glioblastoma cell *in vitro*.

**Supplementary Figure 2.**
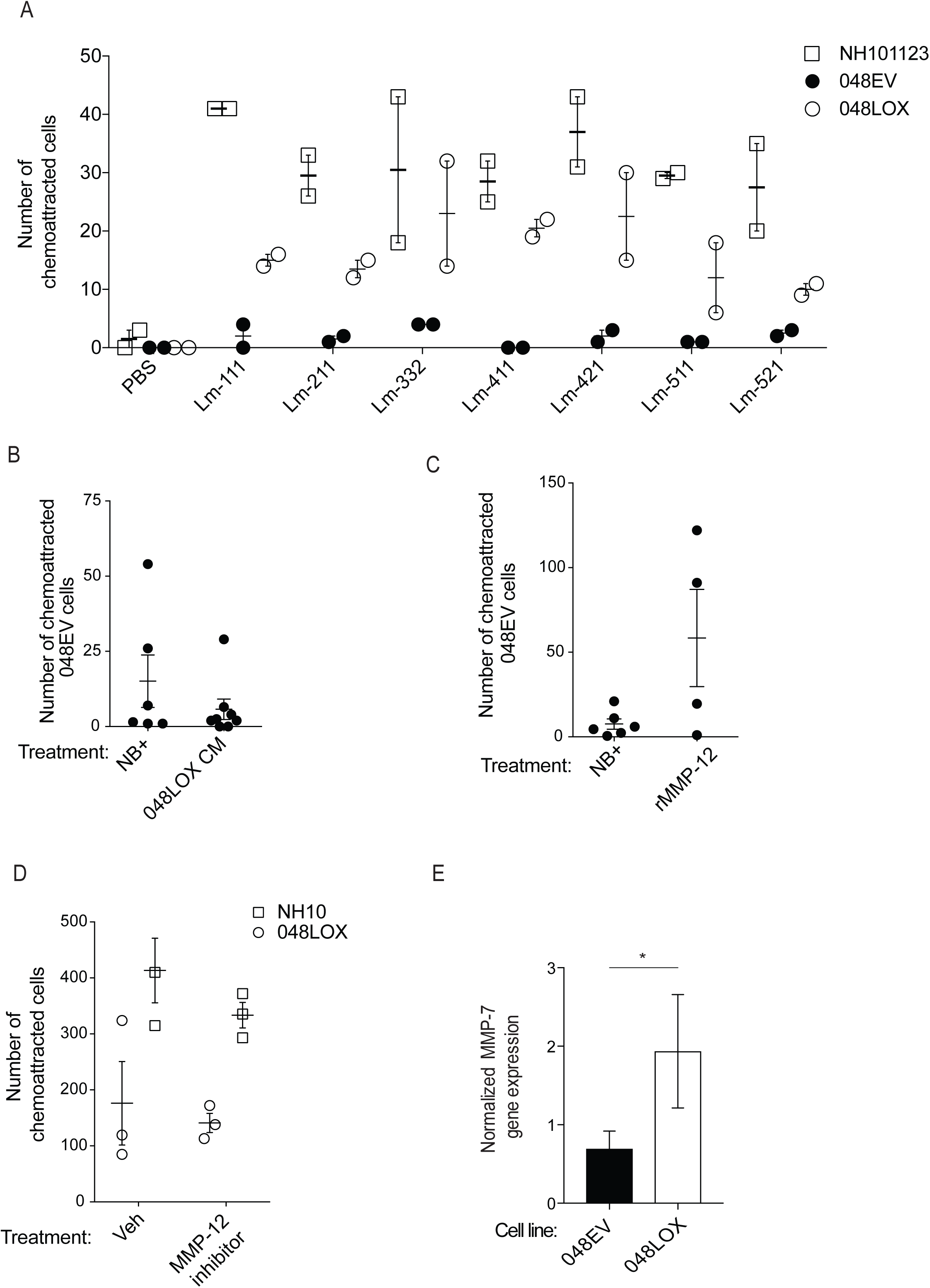
Chemotactic assays.

**Supplementary Figure 3.**
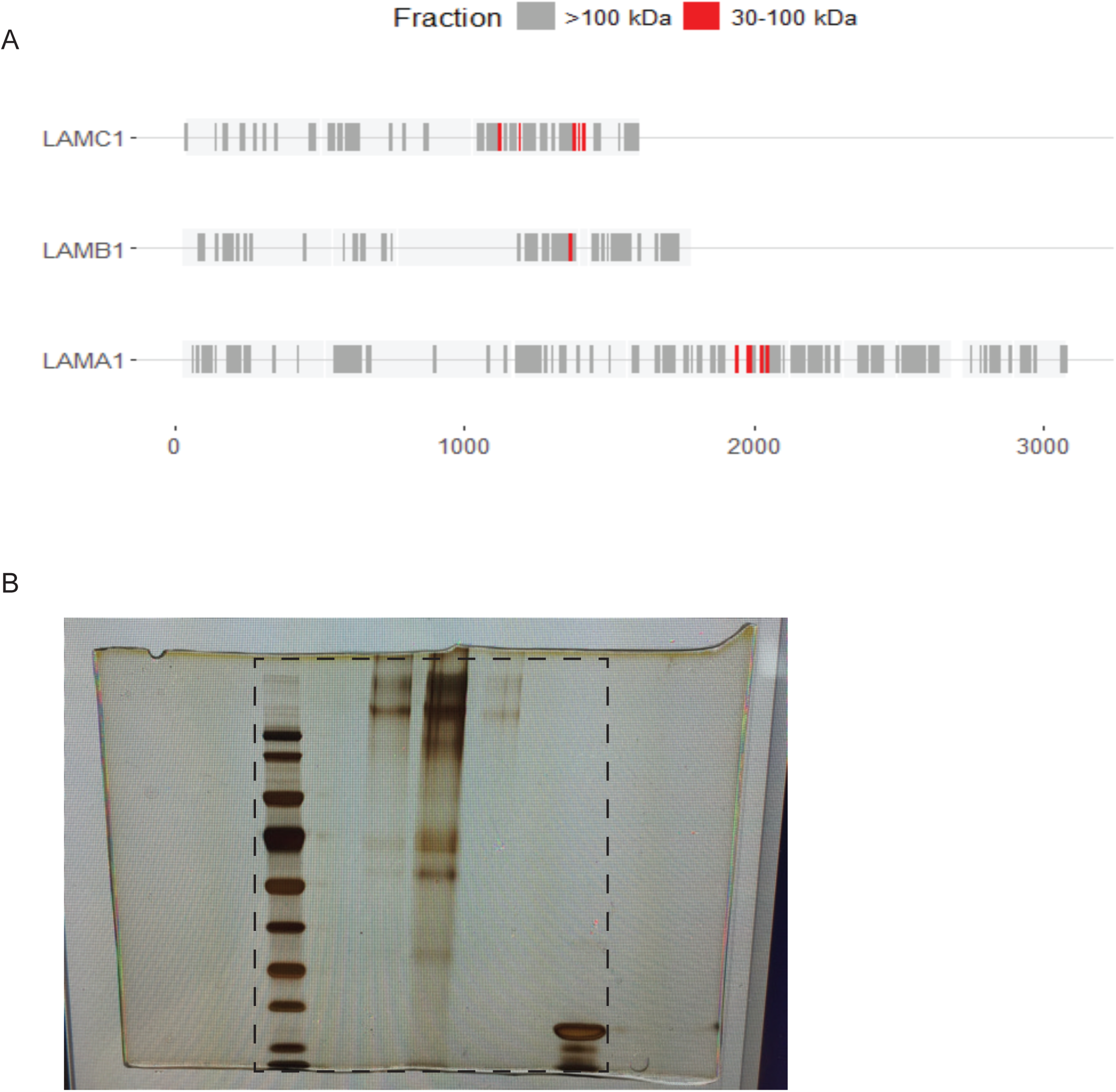
Peptides detected in the MMP-7 - Lm-111 cleavage assay.

